# Tubulin Acetylation Deficiency Promotes Axonemal Turnover and Increases Cytoplasmic Microtubules

**DOI:** 10.1101/2025.10.01.679783

**Authors:** Tomohiro Kubo, Natsumi Tajima-Shirasaki, Rinka Sasaki, Toshiyuki Oda, Masayuki Onishi

## Abstract

Tubulin post-translational modifications regulate microtubule dynamics; among these, α-tubulin acetylation has been linked to microtubule stability. We generated a *Chlamydomonas* mutant lacking the acetyltransferase αTAT1, which completely abolished α-tubulin K40 acetylation. Surprisingly, the lengths of normally acetylated structures, axonemes and rootlets, were largely unaffected. αTAT1 localized to the flagellar tip, suggesting that it is the primary site of acetylation. Loss of acetylation caused an increase in axonemal tubulin turnover, as revealed by dikaryon-fusion assays. Unexpectedly, the *αtat1* mutant displayed an increased number of dynamic cytoplasmic microtubules and could regenerate long flagella after amputation, even when protein synthesis was inhibited. Despite these cytoskeletal changes, steady-state flagellar length, cell growth, and cell division remained essentially normal. These findings suggest that acetylation modulates microtubule behavior by regulating axonemal tubulin turnover and cytoplasmic microtubule dynamics, while cellular morphology is buffered against variations in microtubule content.

## Introduction

Tubulin undergoes several kinds of post-translational modifications (PTMs) such as acetylation, tyrosination, detyrosination, polyglutamylation, polyglycylation and phosphorylation (Westermann and Weber, 2003; Janke and Bulinski, 2011). These PTMs can affect the dynamics or stability of microtubules, as well as the interaction between microtubules and associated proteins. Investigation on the precise function of tubulin PTMs is crucial for understanding the molecular function of microtubules.

Tubulin acetylation, in which an acetyl group is added to the epsilon-amino group of α-tubulin, was discovered in *Chlamydomonas reinhardtii* flagella (L’Hernault and Rosenbaum, 1985). Soon after this discovery, a specific monoclonal antibody against acetylated α-tubulin was produced (Piperno and Fuller, 1985), and the acetylated residue was identified as lysine 40 (K40) (LeDizet and Piperno, 1987). Interestingly, this residue is located on the luminal side of microtubules (Nogales et al., 1999). To investigate the functional importance of tubulin acetylation, mutants expressing non-acetylatable form of α-tubulin, with K40 replaced by another amino acid, were generated in *Chlamydomonas* (Kozminski et al., 1993) and the ciliate *Tetrahymena thermophila* (Gaertig et al., 1995). Both mutants showed no specific phenotype, raising a possibility that α-tubulin acetylation may be non-essential for these organisms.

Later studies with the nematode *Caenorhabditis elegans* identified the enzyme responsible for α-tubulin acetylation, the product of the gene *MEC-17,* and it was termed α-tubulin acetyltransferase 1 (αTAT1) (Akella et al., 2010; Shida et al., 2010). In contrast to the α-tubulin K40 mutations, disruption of *MEC-17* homologues caused microtubule instability in *Tetrahymena* and neuromuscular defects in zebrafish (Akella et al., 2010). In *C. elegans*, mutations in the *MEC-17* gene cause an abnormal function of touch receptor neurons (Chalfie and Au, 1989), which, in the wild-type organism, undergo extensive tubulin acetylation. In mice also, αTAT1 is a major α-tubulin acetyltransferase (Kalebic et al., 2013; Kim et al., 2013); mice lacking αTAT1 are viable and develop normally, but their sperm show abnormal motility and microtubule stability is increased (Kalebic et al., 2013).

Biochemical and structural properties of αTAT1 have been extensively explored. In vitro assays using various types of tubulin polymers demonstrated that αTAT1 can enter the microtubule lumen through open ends or lattice defects (Szyk et al., 2014; Coombes et al., 2016), although it can also bind to the outside surface of the microtubule (Howes et al., 2014). Unexpectedly, despite the luminal localization of K40, its αTAT1-mediated acetylation *in vitro* did not show preference for microtubule ends (Szyk et al., 2014). On the other hand, FRET-based assays demonstrated that tubulin acetylation reduces the nucleation frequency and accelerates shrinkage rates of microtubules (Portran et al., 2017), suggesting that the acetylation critically affects the dynamics of microtubule ends.

In this study, we assessed the *in vivo* function of αTAT1 and α-tubulin K40 acetylation in *Chlamydomonas* by analyzing the phenotype of a mutant, *αtat1(ex3)*, which lacks the sole αTAT1 ortholog in this organism. This mutant completely lacked tubulin acetylation and showed an increase in the turnover of flagellar tubulin and the amount of cytoplasmic microtubules. Despite these alterations, both flagellar length and cell size remained normal. Our observations thus shed light on a novel aspect of acetylation in controlling tubulin dynamics, as well as the cell’s capacity to cope with abnormal cytoskeletal remodeling.

## Results

### Generation of a *Chlamydomonas* mutant lacking αTAT1 by CRISPR/Cas9

In the *Chlamydomonas* genomic database, we found a gene encoding a homologue of mouse α-tubulin acetyltransferase 1 (αTAT1). To explore the role of tubulin acetylation in *Chlamydomonas*, we generated a mutant lacking the αTAT1 activity, using CRISPR/Cas9-mediated gene editing to introduce a hygromycin-resistant cassette into the exon 3 of the *αTAT1* gene (Cre07.g345150) in a wild-type strain (cc124). Genotyping with a specific primer set identified several transformants with a hygromycin-resistant cassette at the target site (Fig. 1 A). Sequencing one such transformant revealed that the insertion created a premature stop codon in the exon 3, suggesting that the transcription of full-length *αTAT1* is blocked. Hereafter, we refer to this allele as *αtat1(ex3)*.

**Figure 1.**
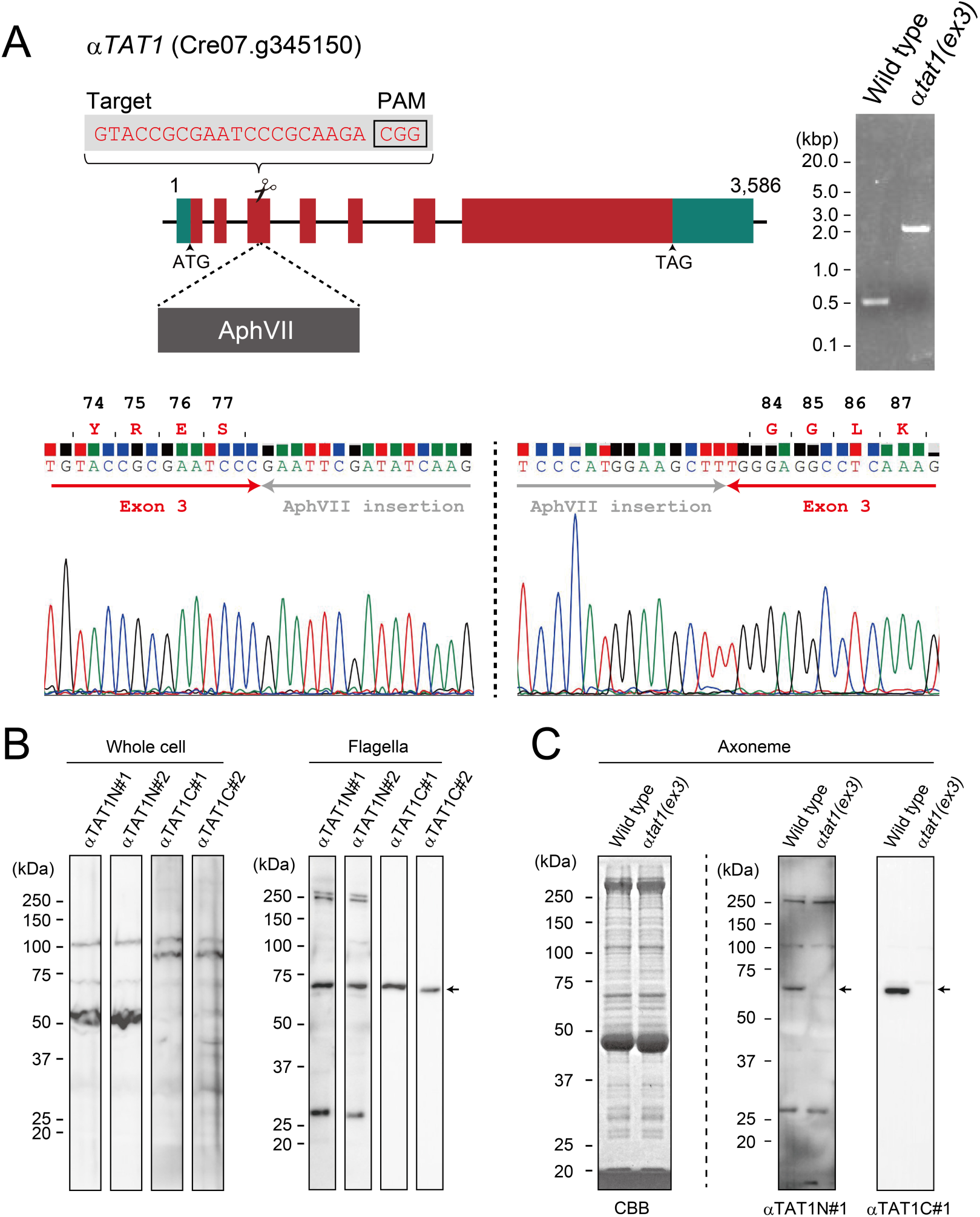
Generation of a *Chlamydomonas* mutant lacking the *αTAT1* gene. (A) The structure of the *αTAT1* (Cre07.g345150) gene. Exons are indicated in dark red, and untranslated regions in greenish blue. As confirmed by gene sequencing and PCR using a specific primer set, a hygromycin-resistant gene cassette (dark gray) was integrated into exon 3 of the *αTAT1* gene. (B) Western blot validation of anti-αTAT1N#1, αTAT1N#2 αTAT1C#1, and αTAT1C#2 antibodies. Whole cell and flagellar samples of the wild-type strain were used. (C) Western blot of the axonemes isolated from wild type and *αtat1(ex3)*. The *αtat1(ex3)* axoneme lacks αTAT1.

To detect αTAT1 protein in the cell, we generated two batches each of antibodies against the N-terminus (amino acids 12-29, anti-αTAT1N#1 and #2) and C-terminus (amino acids 621-636, anti-αTAT1C#1 and #2). Western blot analyses using these antibodies detected only faint αTAT1 signals with several unexpected signals in whole-cell samples of the wild type (Fig. 1 B). These unexpected signals are likely non-specific, detected due to the low abundance of αTAT1. In contrast to whole-cell samples, flagellar and axonemal samples displayed a specific signal of the predicted size of 63.7 kDa with both antibodies (Fig. 1, B and C). This suggests that αTAT1 is concentrated in the flagella. Importantly, no specific signals were detected with either antibody in *αtat1(ex3)* axonemes (Fig. 1 C). The *αtat1(ex3)* axoneme must thus lack any truncated αTAT1, consistent with the total absence of tubulin acetylation in the axoneme of this mutant, as we describe later.

### αTAT1 is the sole enzyme responsible for acetylation of α-tubulin K40 in *Chlamydomonas*

To investigate the level of acetylated α-tubulin in *αtat1(ex3)*, we performed indirect immunofluorescence microscopy (IFM) on the nucleo-flagellar apparatus (NFAp), a cytoskeletal fraction isolated from cell-wall-less cells after detergent treatment (Fig. 2 A; Sanders and Salisbury, 1995). As expected, the wild-type NFAps showed clear signals in both axonemal and rootlet microtubules by staining with the anti-acetylated α-tubulin antibody (Fig. 2 B). However, the NFAps from *αtat1(ex3)* showed no acetylated tubulin signals along the axonemes or along the rootlets derived from the cell body (Fig. 2 B), suggesting that αTAT1 is the only enzyme responsible for α-tubulin K40 acetylation in *Chlamydomonas*.

**Figure 2.**
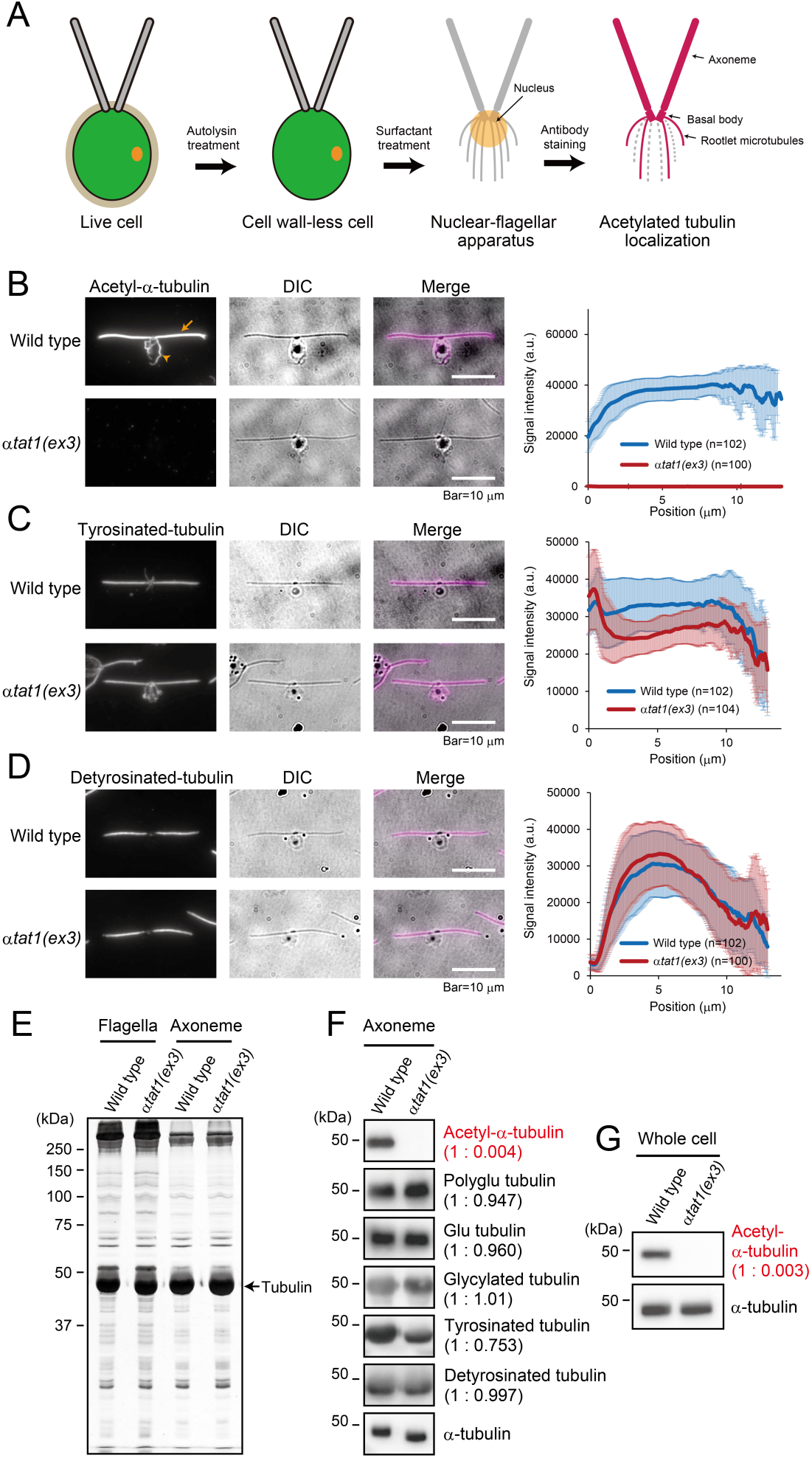
*αtat1(ex3)* completely lacks tubulin acetylation. (A) Diagram of the nuclear-flagellar apparatus (NFAp) sample preparation. Indirect Immunofluorescence microscopy of the NFAps isolated from wild type (cc124) and *αtat1(ex3)* using anti-acetylated α-tubulin (B), tyrosinated tubulin (C), and detyrosinated tubulin (D) antibodies. In (B), an axoneme and a rootlet are indicated by an orange arrow and an orange arrowhead, respectively. Average signal intensities along the axoneme are shown (right panels). (E) CBB-stained SDS-PAGE gel of the isolated flagella and axoneme obtained from wild type and *αtat1(ex3)*. No obvious defects were observed in the bands of *αtat1(ex3)*. Western blot analysis of the axoneme (F) and whole cell extracts (G) comparing wild type (cc124) and *αtat1(ex3)*, using the indicated antibodies. Protein levels were normalized to α-tubulin.

We also examined other types of tubulin modifications in NFAps by IFM. Compared with the wild-type NFAps, polyglutamylation (Fig. S1 A) and glutamylation (Fig. S1 B) in the *αtat1(ex3)* were slightly increased, whereas tyrosination (Fig. 2 C) was slightly decreased in *αtat1(ex3)*. Detyrosination (Fig. 2 D) and glycylation (Fig. S1 C) were almost identical between *αtat1(ex3)* and wild type.

To further confirm these IFM observations, flagella and axonemes were isolated and biochemically analyzed. The protein composition of the *αtat1(ex3)* flagella appeared normal in the SDS-PAGE pattern. Interestingly, the mobility of α-tubulin often appeared slightly higher in *αtat1(ex3)* than in wild type (Fig. 2 E and F). This difference is presumably due to changes in the molecular weight and/or the isoelectric point of α-tubulin resulting from the loss of acetylation. As predicted, the signal of acetylated tubulin was completely missing in *αtat1(ex3)*, both in the axoneme (Fig. 2 F) and in the whole cell (Fig. 2 G). The band shift of total α-tubulin was also observed in the axoneme but not in the whole cell (Fig. 2, F and G), suggesting that the majority of α-tubulin in wild-type is acetylated in the axonemes but not in the cell body. The signal intensity of tyrosinated tubulin in the *αtat1(ex3)* axoneme decreased to approximately 75% of the wild-type level (Fig. 2 F), consistent with the result from IFM observations. This result suggests that tubulin tyrosine ligase, an enzyme responsible for tyrosination of α-tubulin, may interact inefficiently with microtubules lacking acetylated α-tubulin. Western blot signals of other tubulin modifications in *αtat1(ex3)* axonemes were almost identical with those of wild type, confirming the IFM observations. However, unlike the IFM results, no significant differences were found in the signals of polyglutamylated and glutamylated tubulins between wild type and *αtat1(ex3)* (Fig. 2 C); the reason for the apparent discrepancy is unknown. Taken together, these results clearly indicate that the mutant *αtat1(ex3)* completely lacks acetylation of α-tubulin K40 in both flagella and cell body, with a slight decrease in flagellar tubulin tyrosination.

### αTAT1 associates with the axonemal microtubules

To determine the localization and biochemical properties of αTAT1 with high specificity, we generated a strain expressing hemagglutinin (HA)-tagged αTAT1, as the αTAT1 antibodies we raised had low titers and failed to localize it in the axoneme or cell body by IFM (not shown). Using CRISPR/Cas9-mediated gene editing, a sequence encoding triple HA and the 3’ untranslated region (UTR) followed by a hygromycin resistance gene (AphVII) was inserted into a region upstream of the αTAT1’s original 3’UTR in the wild-type strain cc124 (Fig. 3 A), generating *αTAT1-3xHA*. This insertion was expected to produce an αTAT1 lacking the C-terminal five amino acids. IFM of NFAps and Western blotting of isolated axonemes using the anti-acetylated α-tubulin antibody showed that the *αTAT1-3xHA* strain maintains the normal level of α-tubulin K40 acetylation (Fig. 3 B), indicating that the lack of the last five amino acids and the C-terminal tagging have no significant effect on the αTAT1 function. Western-blot analyses of the whole cell and the flagella using an anti-HA antibody detected a single band of the size predicted for the HA-tagged αTAT1 protein (67.2 kDa) in *αTAT1-3xHA* but not in the control cell (Fig. 3 C).

**Figure 3.**
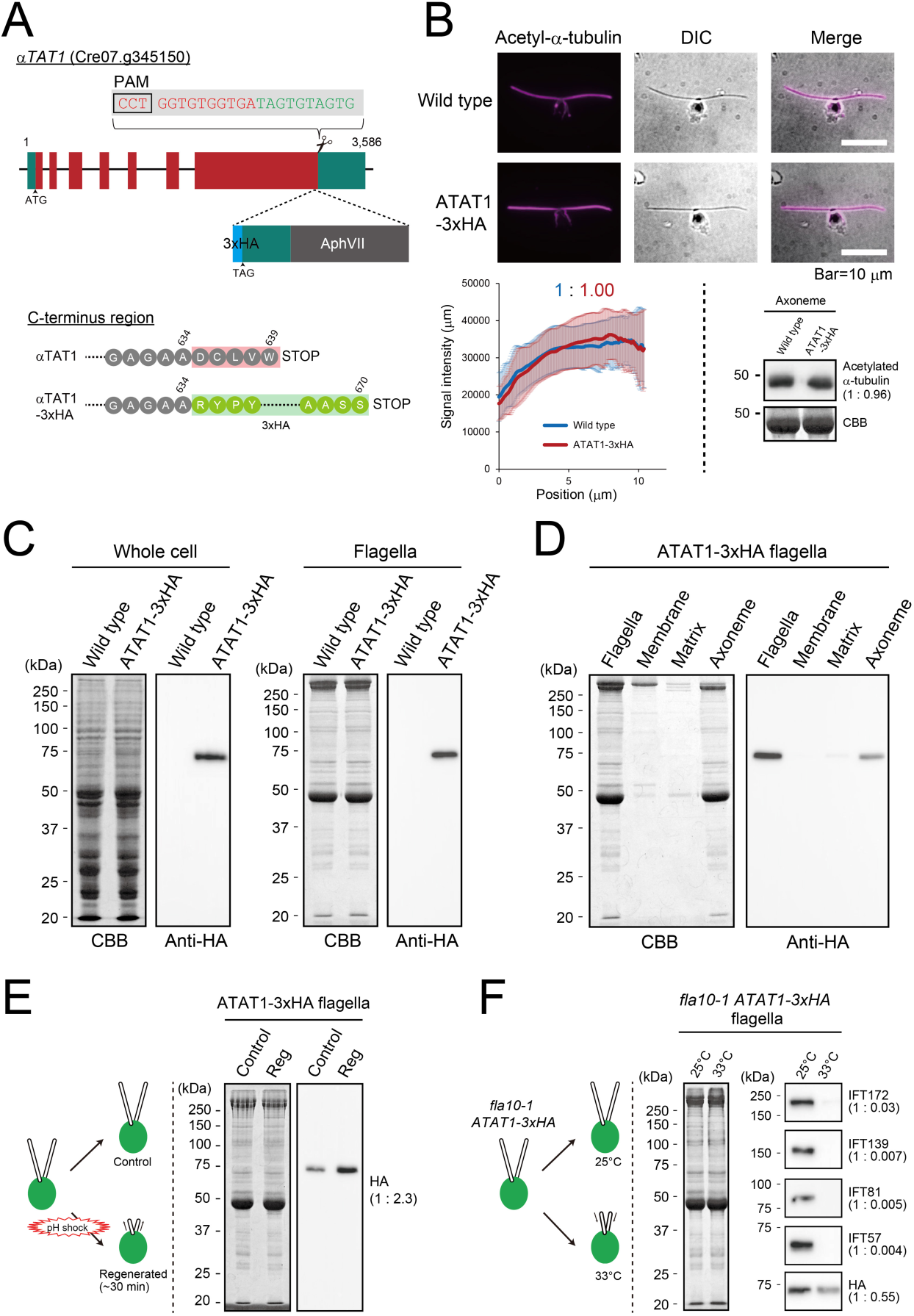
Biochemical properties of ATAT1-3xHA. (A) The structure of the *αTAT1* (Cre07.g345150) gene. The sequence encoding the triple hemagglutinin, 3’UTR, and the hygromycin resistance gene was inserted just before the original 3’ UTR of the *αTAT1* gene of wild type and *fla10-1* strains. The obtained strains were designated *ATAT1-3xHA* and *fla10-1 ATAT1-3xHA*, respectively. (B) Indirect immunofluorescence microscopy of cc124 and the novel *ATAT1-3xHA* strain stained by anti-acetylated α-tubulin. HA-tagging of ATAT1 does not affect its enzymatic activity. (C) Western blot analysis of flagella (right panel) using anti-HA antibody, with the gel stained by CBB. (D) Western blot analysis and CBB-stained gel of flagellar fractions isolated from the *ATAT1-3xHA* strain. ATAT1-3xHA localizes to the axoneme. (E) Western blot analysis and CBB-stained gel of steady-state flagella and regenerated flagella both isolated from the *ATAT1-3xHA* strain. (F) Western blot analysis and CBB-stained gel of flagella isolated from the *fla10-1 ATAT1-3xHA*, incubated at room temperature or at 33°C. Same amount of flagellar samples was loaded and blotted with the indicated antibodies.

Fractionation of *αTAT1-3xHA* flagella revealed that the majority of αTAT1-3xHA was localized to the axoneme, rather than to the flagellar membrane or matrix (Fig. 3 D). This suggests that αTAT1 is associated with the axonemal microtubules to perform α-tubulin acetylation.

To estimate the flagellar amount of αTAT1 during flagellar growth, flagella were isolated from cells 30 min after pH shock-induced flagellar detachment and analyzed by immunoblot (Fig. 3 E). The amount of αTAT1-3xHA per total flagellar protein was approximately 2.3 times higher during flagellar regeneration than in the steady state. Given that flagellar length at 30 min is about half of the steady-state length, this suggests that the total amount of αTAT1 in each regenerating flagellum is similar to that in steady-state flagella.

To examine whether αTAT1 is transported into flagella by intraflagellar transport (IFT), we generated a *fla10-1 αTAT1-3xHA* strain. *fla10-1* is a temperature-sensitive mutant whose IFT stops at 33°C (Kozminski et al., 1995). While clear signals of IFT-B (IFT172, IFT81, and IFT57) and IFT-A (IFT139) complex proteins were detected in the steady-state *fla10-1* flagella isolated at 25°C, none of them were detected in the flagella isolated after the mutant had been incubated at 33°C for 2 hours (Fig. 3 F). After the 33°C incubation, the amount of αTAT1-3xHA in the flagella was decreased to half its original level (Fig. 3 F). This result indicates that the flagellar localization of αTAT1 depends on the IFT.

### αTAT1 localizes to the flagellar tip

The detailed localization of αTAT1 was examined by IFM with an anti-HA antibody on the NFAps of *αTAT1-3xHA*. Specific signals were detected in the *αTAT1-3xHA* NFAps, especially significantly in the distal half of the axoneme (Fig. 4, A and B). The signal intensity sharply increased towards the flagellar tips (Fig. 4, A and B), suggesting that αTAT1 performs tubulin acetylation immediately after tubulins are incorporated into the axoneme. In contrast, no signals were detected in any part of the rootlet microtubules in NFAps (Fig. 4 A), suggesting that the association of αTAT1 with the rootlet might be very small or transient. Flagellar tip localization of αTAT1-3xHA was also observed in the *pf18* mutant lacking the central-pair microtubules (Fig. 4, B and C), although the signal intensity was significantly reduced. The axonemal localization of αTAT1 may partially depend on the presence of the central-pair microtubules or flagellar motility.

**Figure 4.**
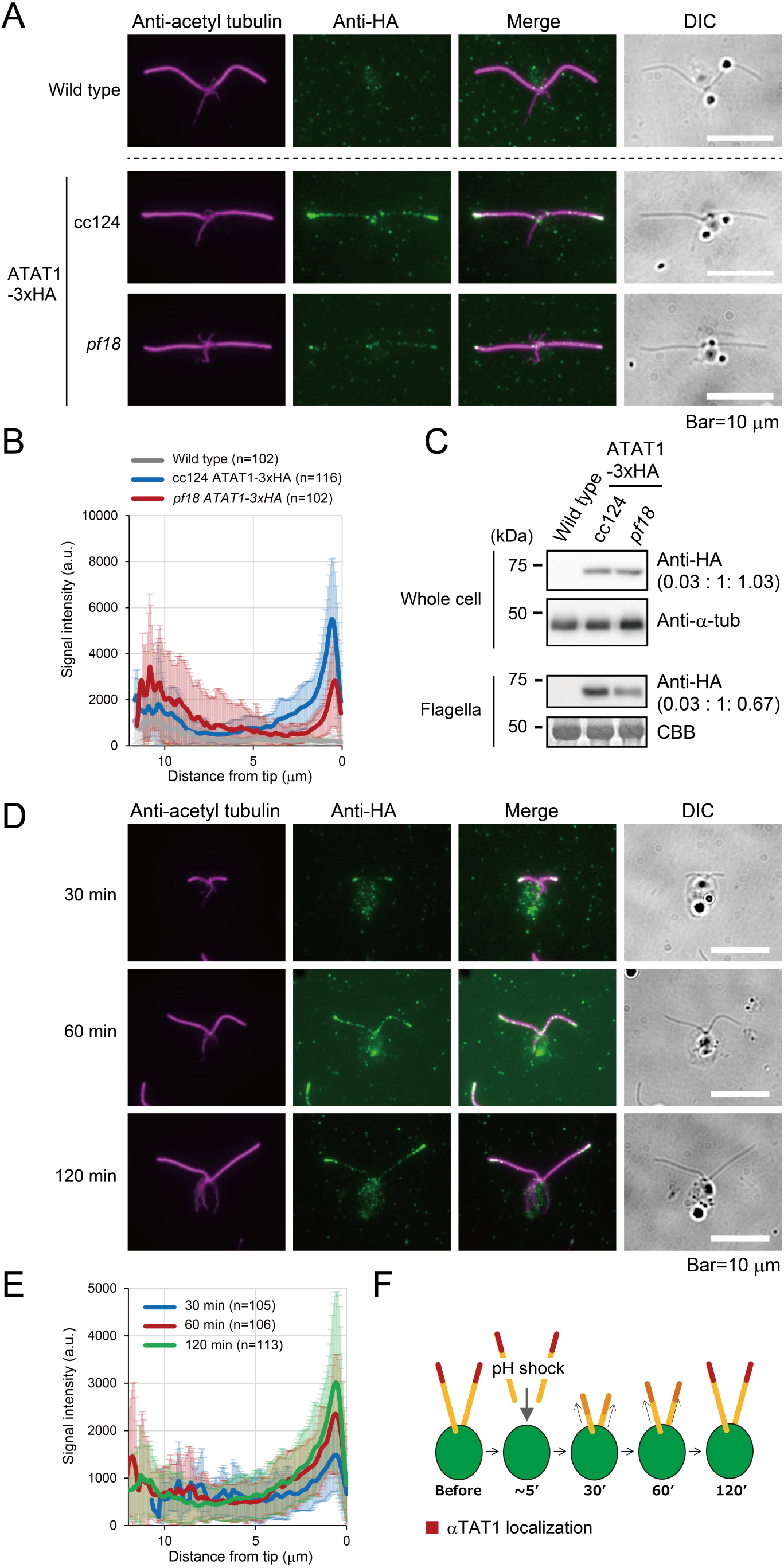
ATAT-3xHA localizes to the flagellar tip. (A) Indirect immunofluorescence microscopy of nuclear-flagellar apparatuses (NFAps) isolated from wild type (cc124), *αTAT1-3xHA*, and *pf18 αTAT1-3xHA* using an anti-HA antibody. (B) Signal intensities from the base to the tip of axonemes were measured in at least 100 cells. The average values and standard deviations are presented as a graph. (C) Western blotting of whole cell and flagellar samples of cc124, *αTAT1-3xHA*, and *pf18 αTAT1-3xHA.* (D) Indirect Immunofluorescence microscopy of the NFAps isolated from wild type (cc124) during flagellar regeneration. Cells were fixed at 30, 60, 120 min after deflagellation, and stained using anti-HA and acetylated α-tubulin antibodies. (E) Signal intensities measured from the axonemal tips are shown. The average values and standard deviations are presented. (F) Schematic illustration of the gradual accumulation of αTAT1 in flagella.

To explore the αTAT1-3xHA localization during flagellar regeneration, we performed IFM 30, 60, 120 min after the αTAT1-3xHA cells were deflagellated by pH shock (Fig. 4 D). Signal intensities from the flagellar tips were measured and averaged in each group (Fig. 4 E). In all groups, the HA signal was most strongly localized at the tip. Additionally, the maximal signal intensity tended to increase with the flagellar length (Fig. 4 E). These results suggest that αTAT1 gradually accumulates around the flagellar tip as flagella elongate, consistent with Western blotting results (Fig. 3 E).

### Tubulin acetylation occurs at the flagellar tip

The tip localization of αTAT1-3xHA raised the possibility that tubulin acetylation first occurs at the flagellar tip. To examine this, we observed dikaryons (Starling and Randall, 1971) formed between wild-type and *αtat1(ex3)* gametes. A dikaryon possesses two pairs of flagella and two nuclei derived from the parental gametes. When a mutant lacking some flagellar components is mated with wild type, the components missing in the mutant are supplied from the shared cytoplasm, and often the deteriorated function of the mutant is “rescued” (Dutcher, 2014). We found that tubulin acetylation started to occur at the flagellar tip within 10 min after mating (Fig. 5 A), and the acetylation signal extended proximally with time (Fig. 5, A and B). These observations suggest that αTAT1 is transported to the tip of *αtat1(ex3)* flagella and carries out α-tubulin acetylation there.

**Figure 5.**
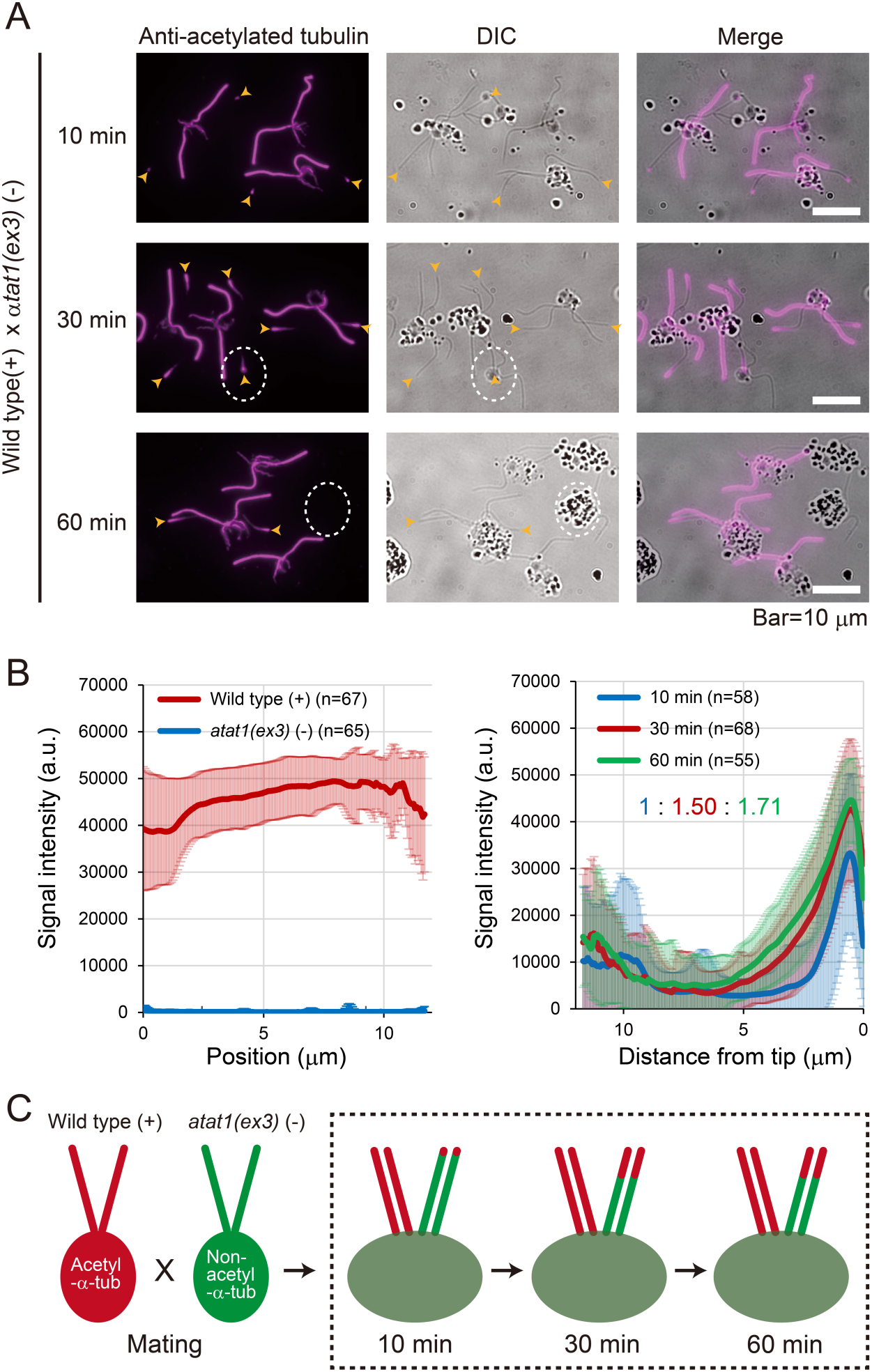
Recovery of α-tubulin acetylation in the dikaryon axonemes. (A) Indirect immunofluorescence microscopy of the NFAps extracted from dikaryons formed between wild type (cc125) and *αtat1(ex3)*, using an anti-acetylated α-tubulin antibody (upper panel). Dikaryons were isolated at 10, 30, and 60 min after mixing the opposite mating types. In the dikaryons, acetylated α-tubulin was newly detected at the tip of the *αtat1(ex3)*-derived flagella (arrowheads). Scattered dots observed in the DIC images are optical artifacts. (B) The average signal intensities of axonemes were compared between cc125 and *αtat1(ex3)* in gametes (left), and among the 10, 30, and 60 min after mixing in dikaryons, focusing only on axonemes derived from *αtat1(ex3)* (right). Intensities along the entire flagella (from tip to base) were averaged, and the mean values were compared. (C) Schematic illustration of this dikaryon rescue.

### Turnover rate of axonemal tubulin is accelerated in the absence of αTAT1

Several studies reported that tubulin acetylation stabilizes microtubules (Cueva et al., 2012; Topalidou et al., 2012; Neumann et al., 2014; Portran et al., 2017; Xu et al., 2017). To examine the stability of axonemal microtubules in the *αtat1(ex3)* mutant, we investigated axonemal tubulin turnover by mating *TUA1-HA* gametes expressing hemagglutinin (HA) tagged α-tubulin (Kozminski et al., 1993) with wild-type gametes. A previous study examining such mating cells showed that HA-tagged α-tubulin becomes incorporated at the tip of the flagella derived from wild type, a phenomenon interpreted as reflecting tubulin turnover (Marshall and Rosenbaum, 2001). IFM and Western blotting using anti-acetylated α-tubulin confirmed that the *αtat1(ex3) TUA1-HA* strain completely lacked tubulin acetylation (Fig. 6, A, B, and C). We then generated two combinations of dikaryons between untagged and *TUA1-HA* gametes: one in the wild-type background [i.e., untagged wild type mated with *TUA1-HA*] and the other in the *αtat1(ex3)* background [i.e., *αtat1(ex3)* mated with *αtat1(ex3) TUA1-HA*]. The dikaryons were fixed and double-stained with anti-HA and tyrosinated tubulin antibodies 10, 30, and 60 min after mating. As reported by Marshall and Rosenbaum (2001), the control dikaryons showed HA-tagged α-tubulin incorporation at the tip of the flagella derived from the wild-type gametes (Fig. 6, C and D). HA-tubulin incorporation was barely detectable at 10 min but became strong at 30 min after mating. In contrast, the dikaryons produced between *αtat1(ex3)* and *αtat1(ex3) TUA1-HA* exhibited more rapid and prominent HA-tubulin incorporation into the flagellar tip. HA signal was already evident 10 min after mating and became even more pronounced in 30 min (Fig. 6, C and D). These results clearly demonstrate that the lack of tubulin acetylation in the axoneme facilitates axonemal tubulin turnover, suggesting that the microtubules in *αtat1(ex3)* flagella are destabilized.

**Figure 6.**
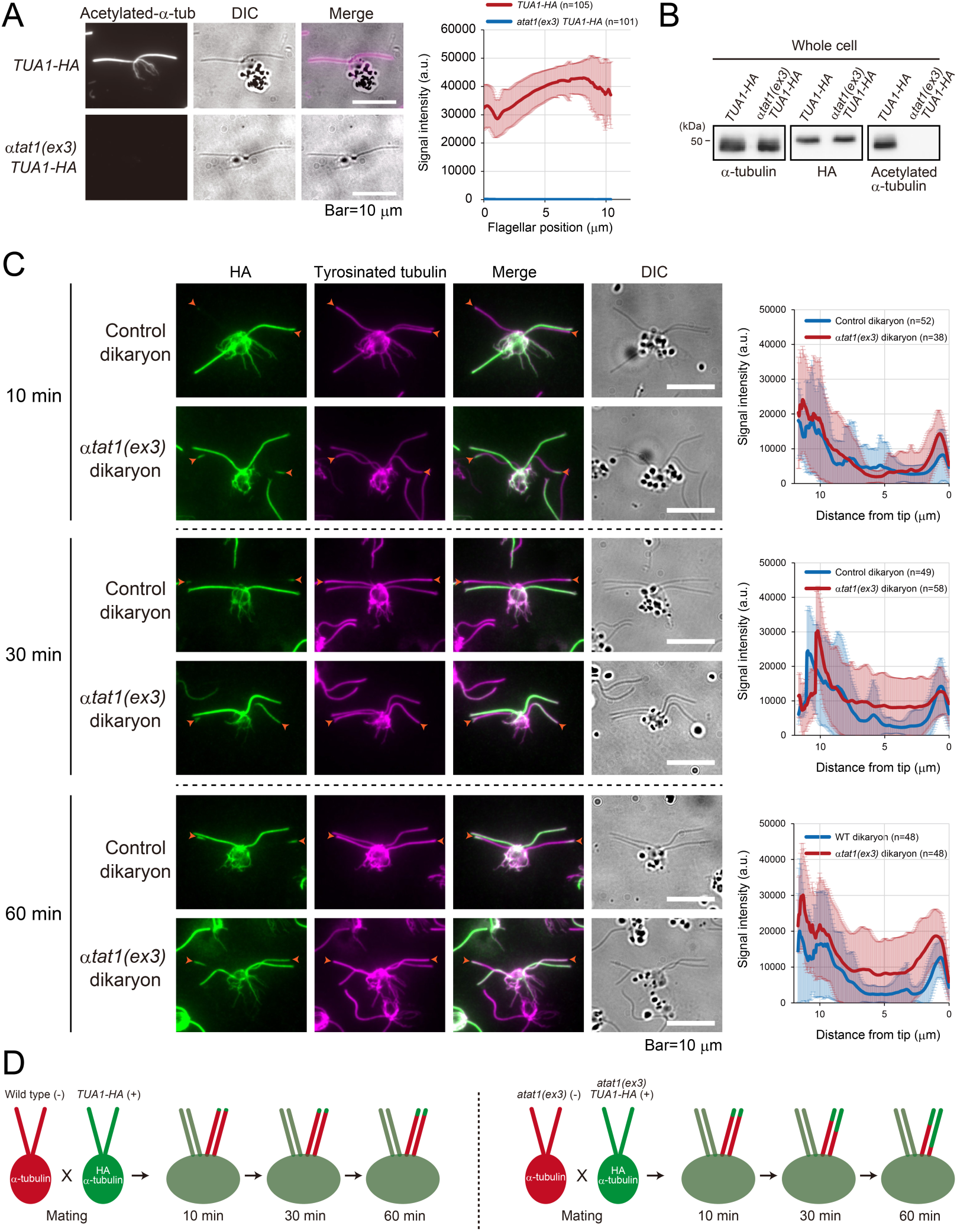
Turnover rate of axonemal tubulin is accelerated in the *αtat1(ex3)* flagella. (A) Indirect immunofluorescence microscopy of the NFAps extracted from *TUA1-HA* and *αtat1(ex3) TUA1-HA*, using anti-acetylated α-tubulin antibody. (B) Western blot analysis of the whole-cell samples using anti-α-tubulin, HA, and acetylated α-tubulin and antibodies. (C) Indirect immunofluorescence microscopy of the NFAps extracted from dikaryons formed between cc124 vs *TUA1-HA*, and *αtat1(ex3)* vs *αtat1(ex3) TUA1-HA*, using an anti-HA and anti-tyrosinated tubulin antibodies. Dikaryons were isolated at 10, 30, and 60 min after mixing the opposite mating types. In the dikaryons, HA tagged α-tubulin was newly detected at the tip of both cc124 and *αtat1(ex3)* derived flagella (arrowheads). (D) Schematic illustration of this dikaryon rescue.

### Loss of tubulin acetylation does not affect rootlet microtubules or cell growth

In wild-type *Chlamydomonas*, the rootlet microtubules in the cell body are also acetylated (Dutcher & O’Toole, 2016) by αTAT1 (Fig. 2, A and B). Tubulin acetylation may well contribute to their intrinsic stability, and the stability may be lost in the *αtat1(ex3)* mutant. We may speculate that the unstable rootlets might in turn affect cell growth and/or cell division. We therefore examined the dynamics of intracellular microtubules by generating wild-type and *αtat1(ex3)* strains expressing superfolder-GFP-tagged α-tubulin (sfGFP-TUA1) (Craft et al., 2015) and culturing them under the 12-h light: 12-h dark cycles for cell-cycle synchronization. Cell sizes in late G1 phase at 8.5, 9.5, and 10.5 hours after the light onset did not show any significant differences between wild-type and *αtat1(ex3)* cells (Fig. 7 A), indicating that the mutation did not significantly affect cell growth. At hour 10.5, many wild-type and *αtat1(ex3)* cells had entered mitosis and cytokinesis with seemingly normal rootlet structures associated with the division site (Fig. 7 B). Quantification of sfGFP-TUA1 signals in the furrow region did not display any difference between the two strains (Fig. 7, C and D). We thus concluded that the *αtat1(ex3)* mutation did not noticeably impair the rootlet structures and stability, or cell growth and division.

**Figure 7.**
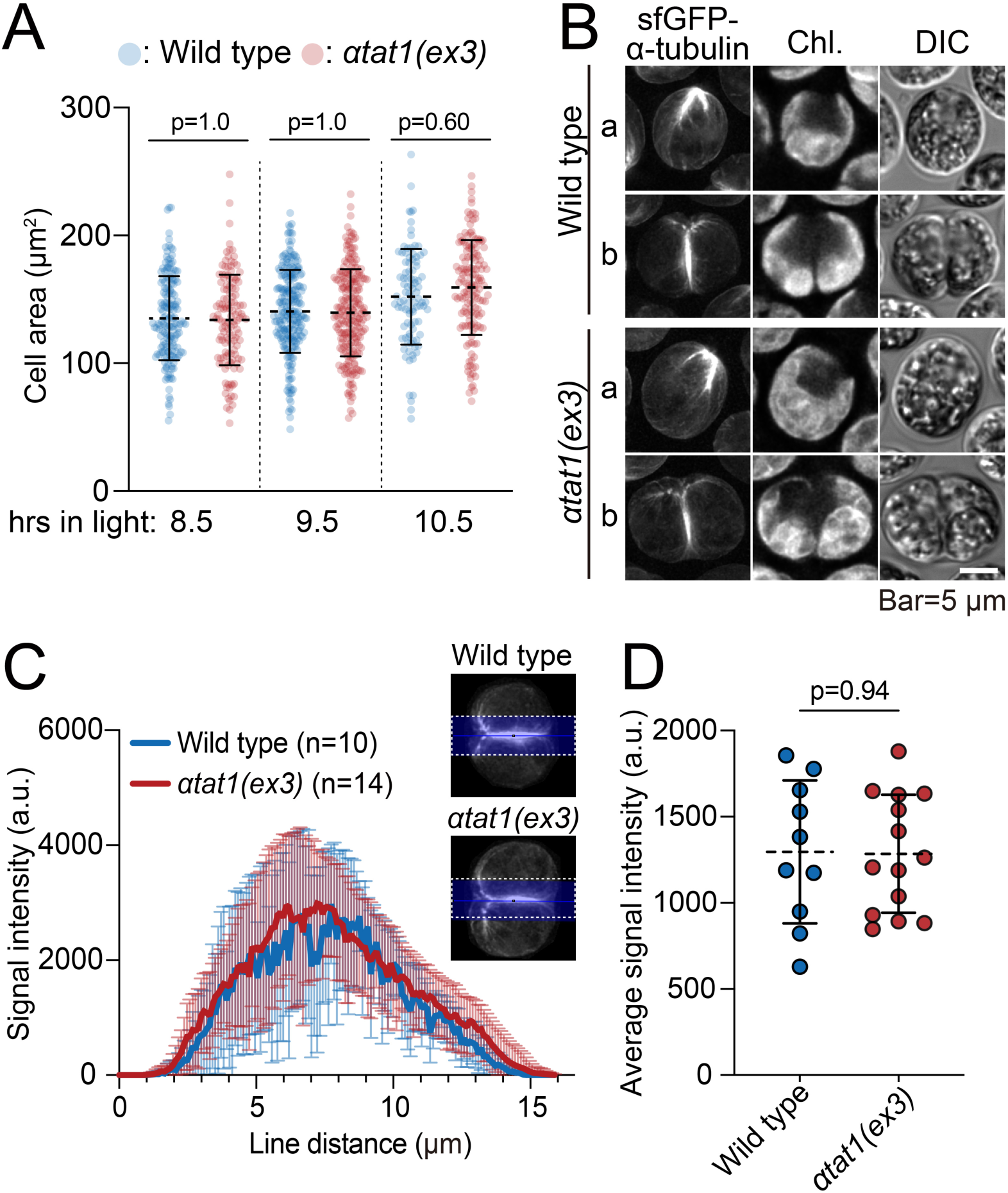
*αtat1(ex3)* shows normal growth, division-site associated rootlets, and cytokinesis. (A) Wild-type and αtat1(ex3) cells expressing sfGFP-TUA1 were cell-cycle synchronized using the 12-h light : 12-h dark cycles. Phase-contrast images of the synchronized populations were taken towards the end of the light phase at hour 8.5, 9.5, and 10.5h, and the cell areas were measured. Values for individual cells (dots) and means (dashed bars) and standard deviations (solid bars) are shown. (B) Cells from hour 10.5 in (A). Representative cells in preprophase (a) and cytokinesis (b) are shown. TUA1-GFP fluorescence, chlorophyll autofluorescence (Chl.), and differential interference contrast (DIC) are shown. (C) Quantification of sfGFP-TUA1 signal intensities along the furrow region during cytokinesis (blue broad line shown in insets). Signal intensities were measured at every 103 nm in 10 wild-type (cc124) cells and 14 αtat1(ex3) cells. The average values and standard deviations are shown. (D) Average sfGFP-TUA1 signal intensities at positions measured in (C) in individual cells.

### The length and number of cytoplasmic microtubules increases in the absence of αTAT1

Besides rootlet microtubules, the *Chlamydomonas* cell body contains 2 to 12 cytoplasmic microtubules that are non-acetylated and highly dynamic (LeDizet and Piperno, 1986; Holmes and Dutcher, 1989; Avasthi and Onishi, 2023; Fig. 2 A). To explore the possible effects of the absence of αTAT1 on the organization of these cytoplasmic microtubules, we stained cytoskeletons isolated from both wild type and *αtat1(ex3)* with anti-α-tubulin antibody. Surprisingly, the cytoplasmic microtubules in these samples clearly differed in the gross appearance: in the *αtat1(ex3)* mutant, they are denser and more widely spread out (Fig. 8 A). Image analysis revealed that both length and number of microtubules were increased (Fig. 8, B and C). This observation was unexpected because tubulin acetylation in the cytoplasm is known to occur only in the stable rootlet microtubules (LeDizet and Piperno, 1986).

**Figure 8.**
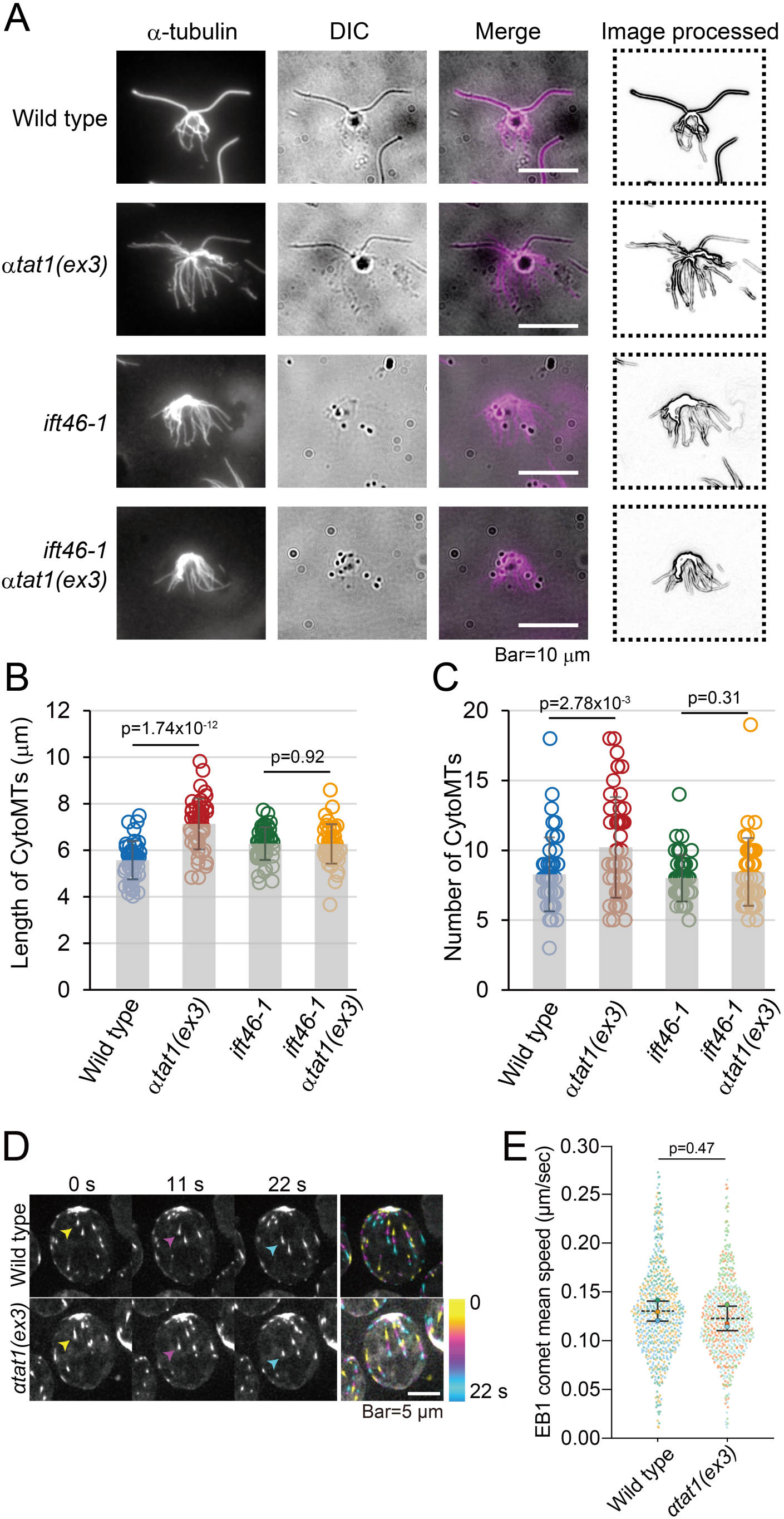
Cytoplasmic microtubules are increased in *αtat1(ex3)* (A) Indirect Immunofluorescence microscopy of the NFAps isolated from wild type (cc124) *αtat1(ex3)*, *ift46-1*, and *ift46-1 αtat1(ex3)* using anti-α-tubulin antibody. The edges of the structures were detected using ImageJ, and the processed images were used to analyze the lengths (B) and the numbers (C) of the cytoplasmic microtubules. (D) Time-lapse images of EB1-mNeonGreen comets in wild-type and *αtat1(ex3)* cells. Representative comets are highlighted with arrowheads at different time points (0 s, 11 s, and 22 s). The rightmost panels show time-color-coded overlays of EB1-mNeonGreen comet trajectories over 22 s. The full series is presented in Movie S1 for wild type and Movie S2 for *αtat1(ex3)*. (E) Scatter plot of EB1 comet mean speeds (µm/sec) in wild-type and *αtat1(ex3)* cells. Each small dot represents an individual comet, with three biological replicates shown in different colors for both wild-type and *αtat1(ex3)*. Large dots, means for each biological replicate. Dotted and solid lines, means ± standard deviations of three replicates.

Speculating that the increase in cytoplasmic microtubule in *αtat1(ex3)* was somehow related to the presence of flagella, we examined the cytoskeleton isolated from the flagella-lacking *ift46-1* mutant (Hou et al., 2007) with or without the *αtat1(ex3)* mutation. Strikingly, no difference was observed between the cytoplasmic microtubules from the *ift46αtat1(ex3)* mutant and those from *ift46* (Fig. 8, A, B, and C). Thus, the increase of cytoplasmic microtubules in the *αtat1(ex3)* mutant indeed appears to depend on the presence of flagella.

If the increased lengths and numbers of cytoplasmic microtubules could act as a buffer to accommodate the excess free tubulin, the concentration of free tubulin available to the growing tip of individual cytoplasmic microtubules should be equal between wild type and *αtat1(ex3)*. Because the microtubule growth rates generally correlate with free tubulin concentrations (Walker et al., 1988), we performed time-lapse microscopy of the microtubule plus-end binding protein EB1 tagged with mNeonGreen (Harris et al., 2016). A strain expressing EB1-mNeonGreen (EB1-NG) was crossed with *αtat1(ex3)*, and the EB1 fluorescence was tracked in progenies with or without *αtat1(ex3)* using three independent sets of segregants (Fig. 8 D). However, no significant difference was observed between wild type and *αtat1(ex3)*: the velocities of the EB1-NG movement were 0.13 ± 0.1 µm/sec in wild type and 0.12 ± 0.1 µm/sec in *αtat1(ex3)* (Fig. 8 E). The lack of αTAT1 thus does not appear to affect the growth rate of cytoplasmic microtubules, confirming the above hypothesis.

### The *αtat1(ex3)* mutant possesses an excess amount of flagellar precursor

After flagellar amputation, *Chlamydomonas* cells can regenerate full-length flagella within 2 hours. Even after the protein synthesis is blocked, they can still regenerate half-length flagella (~6 mm) (Rosenbaum et al., 1969; Lefebvre et al., 1978). The production of short flagella in the presence of a protein-synthesis inhibitor indicates the presence of a certain amount of ‘flagellar precursor’ within the cytoplasm (Rosenbaum et al., 1969; Lefebvre et al., 1978). The main component of the flagellar precursor is thought to be tubulin, stored either as unpolymerized tubulin or as cytoplasmic microtubules (Wang et al., 2013).

To investigate the amount of the flagellar precursor in *αtat1(ex3)*, we performed a flagellar regeneration assay with or without the protein-synthesis inhibitor, cycloheximide. As expected, in the absence of cycloheximide, both the *αtat1(ex3)* and wild-type cells regenerated normal-length flagella within 120 min (Fig. 9 A). When treated with cycloheximide, wild-type cells regained flagella of only half the normal length, consistent with the previous report (Rosenbaum et al., 1969). However, *αtat1(ex3)* cells regenerated significantly longer flagella under the same conditions (Fig. 9 A), suggesting that the mutant possesses an increased amount of flagellar precursor.

**Figure 9.**
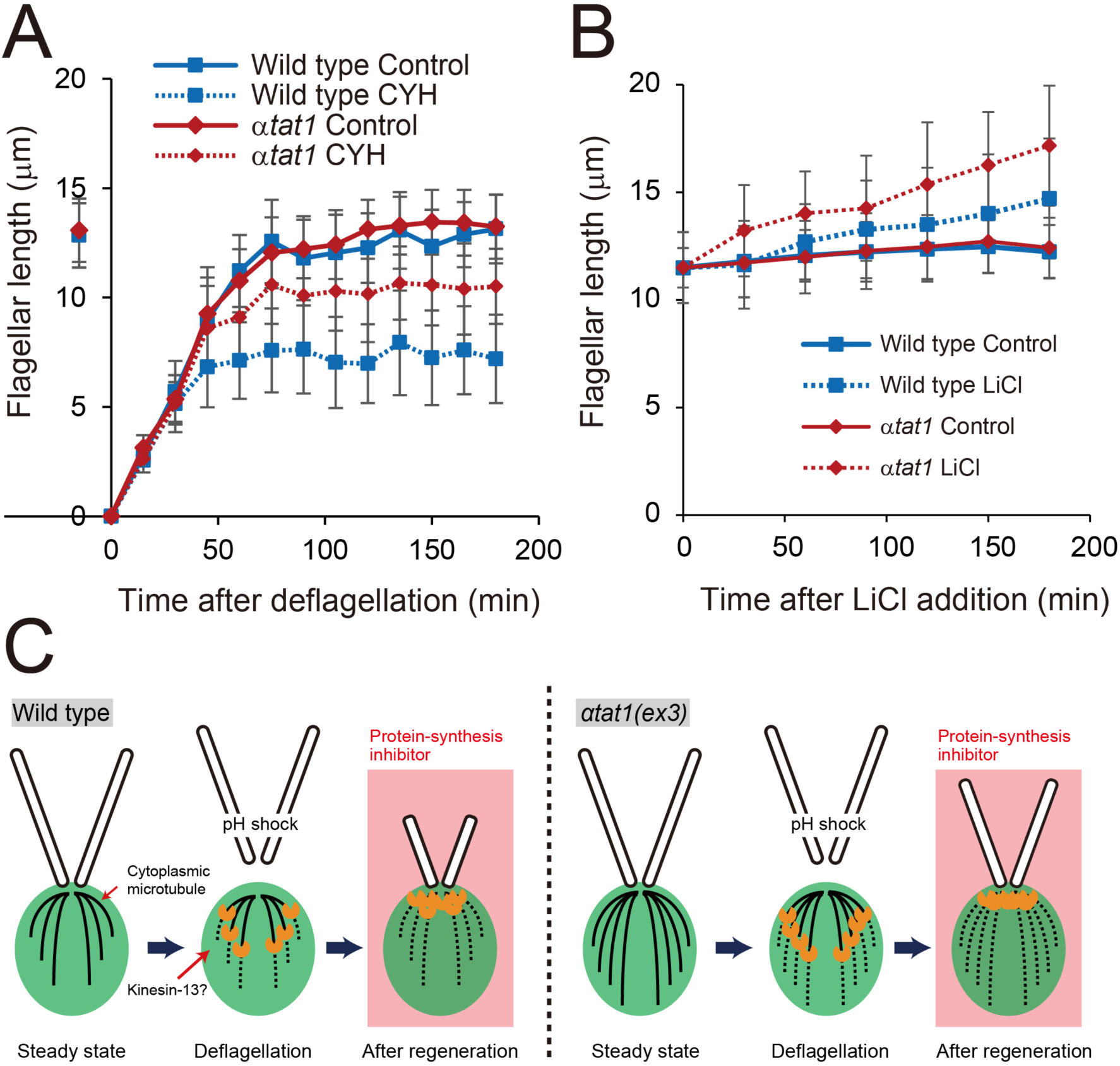
Flagellar precursor is abnormally increased in *αtat1(ex3)* (A) Flagellar regeneration kinetics with and without 30 µg/ml cycloheximide. (B) Flagellar elongation induced by 25 mM LiCl. (C) Schematic illustration of the flagellar regeneration in the presence of protein-synthesis inhibitors. Because *αtat1(ex3)* has an increased amount of cytoplasmic microtubules available for generating flagella, it is able to regenerate longer flagella than the wild type in the presence of protein synthesis inhibitor.

In another line of experiments, we examined the elongation of flagella in the presence of LiCl (Nakamura et al., 1987), a phenomenon that also reflects the amount of flagellar precursor in the cytoplasm (Wilson and Lefebvre, 2004). We found that, after treatment with 25 mM LiCl for 180 min, the flagella in *αtat1(ex3)* elongated more significantly than in wild type (Fig. 9 B). This result further confirms the increased availability of flagellar precursors (including assembly-competent tubulin) in *αtat1(ex3)* (Fig. 9 C).

## Discussion

### The *αtat1(ex3)* mutant completely lacks α-tubulin acetylation

The complete absence of α-tubulin K40 acetylation in *αtat1(ex3)* mutant indicates that αTAT1 is the sole enzyme for this modification in *Chlamydomonas*. However, we cannot tell whether αTAT1 acetylates only α-tubulin K40. To understand the specific effect of α-tubulin K40 acetylation, it may be necessary to analyze mutants in which the endogenous α-tubulin K40 is substituted with other amino acids. We are currently working on this project. We also found that the *αtat1(ex3)* axonemes underwent slightly lower tubulin tyrosination. Tubulin tyrosine ligase may bind with the α-tubulin C-terminus less efficiently when its K40 acetylation is lacking. Furthermore, IFM indicated an apparent increase in tubulin glutamylation and polyglutamylation, whereas Western blotting detected no significant changes in the levels of these modifications in the *αtat1(ex3)* axoneme compared with wild type. We speculate that the lack of α-tubulin K40 acetylation may subtly alter axonemal architecture, thereby changing the accessibility of glutamylated epitopes to antibodies in IFM. Further experiments will be required to clarify the overall alterations in tubulin modifications in the *αtat1(ex3)* axoneme.

### αTAT1 acetylates tubulin at the flagellar tip

In mammals, αTAT1 has been shown to localize along the length of various types of cilia, such as the motile cilia in the brain, the primary cilia in the kidney, and the connecting cilia in the retina (Nakamura et al., 2016). In constrast, *Chlamydomonas* αTAT1 localizes at the tips of flagella, where most α-tubulin acetylation likely takes place (Fig. 4 A). This idea is supported by the recovery of acetylated tubulin signals at *αtat1(ex3)* flagellar tips in dikaryon cells formed between wild type and *αtat1(ex3)* (Fig. 5 A). The flagellar tip localization of the newly acetylated α-tubulin and αTAT1 highlights this region as a critical site for the regulation of the αTAT1 activity as well as the flagellar microtubule dynamics. Because *Chlamydomonas* cells undergo flagella resorption and regeneration once every cell-division cycle, tubulin acetylation at the growing tip during regeneration may be sufficient for this tubulin code to be retained until next division under normal growth conditions.

Despite the observed tip localization of αTAT1, *in vitro* studies have shown that it can bind to microtubules uniformly along their length, independent of the distance from the plus and minus ends (Howes et al., 2014). This suggests that some factors beyond the microtubule itself, such as the IFT machinery or tip-associated scaffolds, may be responsible for directing αTAT1 to the flagellar tip *in vivo*. However, we have not been able to identify any αTAT1-interacting proteins by immunoprecipitation using αTAT1-3xHA flagella (not shown). The question of whether the tip localization of αTAT1 is conserved in organisms other than *Chlamydomonas* also remains to be investigated.

### Possible mechanisms of rootlet microtubule acetylation

Among the cytoplasmic microtubules of *Chlamydomonas*, only four stable bundles called rootlet microtubules undergo acetylation (LeDizet and Piperno, 1986). Since our antibody did not detect cytoplasmic αTAT1 signals in Western blots of cell bodies or in the IFM images of NFAps of wild type, the amount of αTAT1 in the cytoplasm could be very low. There may be two possible mechanisms that produce acetylated rootlet microtubules; one is that acetylation is carried out in the cytoplasm by the undetectably small amount of cytoplasmic αTAT1; the other is that acetylation occurs exclusively in the flagella and that the rootlet microtubules incorporate the acetylated tubulin derived from axonemal microtubules, which undergo depolymerization during the flagellar resorption in late G1 phase (Cavalier-Smith, 1974). Further studies are required to understand how rootlet microtubules become specifically acetylated. Related to this problem, how deacetylation enzymes such as HDAC-6 (Zhang et al., 2003) and SIRT-2 (North et al., 2003) choose the right microtubules to work on also remains to be investigated.

### α-tubulin acetylation as a regulator of microtubule dynamics

The lack of acetylation significantly increased the turnover rate of axonemal tubulin, as revealed by the dikaryon fusion assay (Fig. 6, C and D). This finding appears to be in line with the previously proposed idea that acetylation contributes to the stabilization of axonemal microtubules (LeDizet and Piperno, 1986; Shida et al., 2010). Surprisingly, cytoplasmic microtubules were increased in the *αtat1(ex3)* mutant (Fig. 8 A). One possible hypothesis is that the accelerated axonemal turnover releases excess tubulin dimers, which are subsequently captured and reutilized for the assembly of cytoplasmic microtubules. This model suggests that axonemal and cytoplasmic microtubules are not independent but share a common pool of tubulin as previously suggested (Wang et al., 2013), and that perturbations in axonemal dynamics can influence cytoplasmic microtubule organization. This hypothesis is consistent with the observation that expansion of cytoplasmic microtubules did not occur when flagella were absent, even in the background of *αtat1(ex3)* mutation (Fig. 8, A to C). Another possible hypothesis assumes no direct link between axonemal turnover and the quantity of cytoplasmic microtubules. Instead, the loss of tubulin acetylation may affect the overall tubulin protein level, leading to an accumulation of excess tubulin that is subsequently used for the assembly of cytoplasmic microtubules. This could occur through reduced degradation or altered recycling of tubulin, which together lead to its accumulation in the cytoplasm. It is noteworthy that despite the significant change in microtubule dynamics and quantity in *αtat1(ex3)*, both flagellar length and cell size remained unchanged. Cells can thus tolerate such changes without altering overall morphology, a phenomenon that also warrants further investigation.

## Materials and Methods

### Strains and Cultures

The strains used in this study are listed in Supplemental Table 1. For long-term maintenance, the cells were cultured on Tris-acetate-phosphate (TAP; Gorman and Levine, 1965) agar plates at 15°C. For the experiments, the cells were proliferated and cultured in TAP liquid medium at 25°C with a 12:12 hour light/dark cycle. Cell-cycle synchronization was done on TAP agar using 12:12 hour light/dark cycles as described in Onishi et al. (2020). The genome editing experiments to generate *αtat1(ex3)* and *αTAT1-3xHA* mutants were performed according to Kubo et al. (2023). The crRNAs are listed in Supplemental Table 2. Genetic transformation to create sfGFP-TUA1 (Craft et al., 2015) and EB1-mNeonGreen (Harris et al., 2016) strains was done by electroporation using CHES buffer and a NEPA21 as described in Onishi et al. (2016).

### Assessment of swimming velocity

Swimming velocity was acquired by tracking images of the moving cells. Briefly, the cells under the dark-field microscope equipped with 40x objective were recorded using a digital camera with a frame rate of 30 fps, and the obtained movies were processed with ImageJ.

### Measurements of flagellar regeneration kinetics

To measure flagellar regeneration kinetics, cells were first deflagellated by adjusting the pH of the liquid culture to 4.5 using 0.5 M acetic acid. After a 20-second incubation at pH=4.5, the pH was neutralized to 7.4 with 0.5 M KOH. Aliquots of the cells were collected at 15-minute intervals (up to 180 min) and fixed with 0.3% glutaraldehyde. To obtain the average flagellar length, at least 50 images of flagella were traced and measured using ImageJ.

### Indirect immunofluorescence microscopy

Immunostaining of nuclear-flagellar apparatuses (NFAps) was carried out following Kubo et al. (2024). Fully grown cells were suspended in 6 ml of handmade autolysin (Picariello et al., 2020) and gently agitated for 60 min to remove cell walls. The cells were washed with NB buffer [6.7 mM Tris-HCl (pH 7.2), 3.7 mM EGTA, 10 mM MgCl_2_, and 0.25 mM KCl] and placed on an 8-well slide glass (8 mm well; Matsunami) treated with polyethylenimine. The cells were demembranated with 1% Igepal CA-630 (Sigma) and subsequently fixed with 2% paraformaldehyde in NB buffer for 10 minutes. The resulting NFAps were treated with acetone and then methanol, both at −20°C for 5 minutes. After rehydration by 0.05% Triton X100 in PBS, the NFAps were treated with blocking buffer (1% BSA and 3% Fish Skin gelatin in PBS) and incubated with primary antibodies (Supplemental Figure 3) followed by secondary antibodies (goat anti-rabbit IgG Alexa 488, 1:200, Invitrogen; goat anti-mouse IgG Alexa Fluor 594, 1:200, Invitrogen; goat anti-rat IgG Alexa Fluor 488, 1:200). The NFAps were treated with antifade mountant (SlowFade Diamond, Thermo Fisher Scientific) and encapsulated with a glass cover slip. Imaging was done using a microscope (BX53, Olympus) equipped with a 100× UPlan FL N Oil objective lens (Olympus) and a ORCA-Flash4.0 sCMOS camera (Hamamatsu).

For the immunofluorescence of dikaryon cells, the following procedures were carried out before the autolysin treatment. Cells of opposite mating types were cultured on a TAP-agar plate for 6 to 7 days. The cells were then suspended in M-N liquid medium and gently shaken for three hours to induce gametogenesis (Sager and Granick, 1954). The gametes were subsequently treated with autolysin for an hour to remove cell walls. The mating type plus and minus gametes were mixed and incubated for various periods. The subsequent processes of cell fixation and antibody treatments were carried out as described above.

### Isolation of flagella

Flagella were isolated according to Witman et al. (1972) with some modifications. Basically, the purification process was carried out at 4°C. Two liters of fully grown cells were collected with centrifugation (3,000 rpm, 5 min, Hitachi CR21N, angle rotor R10A3) and washed with 30 ml of 10 mM HEPES (Nakalai tesque). The cells were then suspended in 10 ml of HMS buffer (10 mM HEPES, 5 mM MgSO_4_, 4% sucrose) and treated with 1.6 ml of 0.1 M dibucaine-HCl (Wako) to detach flagella. 10 ml of HMS buffer with 1 mM EGTA was added to the deflagellated cells. The cell suspension was centrifuged twice to remove cell bodies; after the first centrifugation (3,000 rpm, 3 min, Himac CF18R, swing rotor T4SS31), the supernatant was collected and centrifuged again under the same conditions. To achieve further purity of the flagella, a sucrose cushion centrifugation was occasionally applied (Craige et al., 2013). The supernatant was then centrifuged to collect flagella (15,000 rpm, 20 min, Himac CF18R, angle rotor T15A41). The flagellar pellet was suspended in HMDEK buffer (30 mM HEPES, 5 mM MgSO_4_, 1 mM DTT, 0.1 mM EGTA, and 25 mM CH3COOK). To obtain axonemal samples, the flagella were demembranated using 0.1% Igepal CA-630 (Sigma), washed once (15,000 rpm, 10 min, Himac CF16RX, Hitachi T16A31 rotor), and resuspended in HMDEK buffer. After measuring their concentration using a spectrophotometer (ASONE ASD11D), flagellar or axonemal samples were mixed with handmade 4xSDS sample buffer and boiled for 5 min to prepare 1 mg/ml SDS-PAGE protein samples.

### Preparation of whole cell samples

Two different methods were employed to prepare whole-cell samples for Western blotting. For the analysis of acetylated tubulin, 10 ml of fully grown cell culture was centrifuged at 3,000 rpm for 3 min at room temperature. The resulting cell pellet was resuspended in 50 µl of HMDEK buffer, frozen in liquid nitrogen, and thawed twice. The lysate was then centrifuged at 16,000 rpm for 5 min at 4°C. The protein concentration of the supernatant was measured, and the sample was adjusted to 1 mg/ml by adding HMDEK buffer and 4xSDS sample buffer. The signal from the anti-α-tubulin antibody was used as a loading control. For the analysis of total cellular tubulin levels, 10 ml of cell culture at a density of 1×10^6^ cells/ml was centrifuged at 3,000 rpm for 3 min at room temperature. The pellet was resuspended in 480 µl of 1xSDS sample buffer and boiled for 5 min.

### Western blotting

Protein samples were subjected to electrophoresis using a handmade SDS-PAGE gel containing 9% acrylamide (Nakalai Tesque) and transferred onto a PVDF membrane (Immobilon-P, size 0.45 µm; Merck-Millipore). The membrane was first treated with blocking buffer [5% skim milk in PBST (Phosphate-Buffered Saline with 0.1% Tween 20)]. Subsequently, the membrane was probed with primary antibodies (Supplemental Table 3) in the blocking buffer for 45 minutes and washed three times with the blocking buffer for 5 minutes each. The membrane was then probed with secondary antibodies (goat anti-mouse IgG (H+L) (Invitrogen, Cat#31430) or goat anti-rabbit IgG (H+L) (Invitrogen, Cat#31460)) for 45 min and washed three times with PBST for 5 min each. The membrane was treated with Chemi-Lumi One Super (Nakalai Tesque, #02230) and visualized using the Fusion Solo S system (Vilver Bio Imaging).

### Generation of antibodies

The antibodies against αTAT1 and RPL4 (Cre09.g397697) were produced and purified by Bio-Synthesis (https://www.biosyn.com/). The αTAT1 peptides used to immunize rabbits were GEQHISFWDSKRIATLKP-Cys (N-terminus) and KLMAVKQRSGAGAAD-Cys (C-terminus). For RPL4, the immunizing peptide was Cys-KNMLVESDYAGDDYD (C-terminus). The powdered antibodies were reconstituted with the specified amount of water and subsequently used for the experiments.

### Live-cell microscopy

Still images were acquired using a Leica THUNDER 3D Cell Culture Imager equipped with an HC PL Apochromat 63X/1.40 oil-immersion objective lens and a Leica DFC9000 GTCVSC-10038 camera. Images were captured using Leica Application Suite (LAS) X software. GFP- and mNeonGreen-tagged proteins were detected using a 510 nm excitation LED and 535/15 nm emission filters. Chlorophyll autofluorescence was captured using 640 nm excitation and 705/72 nm emission. Prior to imaging, cells were scraped from the agar surface and placed on a block of TAP + 1.5% low-melting-point agarose (IBI Scientific, CAT# 89125-532). To enhance image clarity, Thunder Large Volume Computational Clearing was applied at 92% strength with a 1000-nm feature scale. Image post-processing, including intensity adjustments and quantitative analysis, was performed using ImageJ (National Institutes of Health). All images from a single experiment with the same strain were processed under identical settings to ensure comparability across figures.

### EB1 tracking

EB1-mNG comets were imaged for 5 min with 1-second exposure and 11-second intervals. Comet tracking was performed using TrackMate v7.12.2 (Tinevez et al., 2017) in Fiji (ImageJ). Spots were detected with the LoG (Laplacian of Gaussian) detector with an estimated object diameter of 0.7 µm and an initial quality threshold of 10. No median filter was applied, and subpixel localization was enabled to enhance detection accuracy. For comet tracking, the LAP tracker was used with a maximum search radius of 3 µm and gap closing disabled. To refine the analysis, filters were applied to the tracks; only comets with a total travel distance above 1.38 µm and a mean directional change rate below 0.12 radians per second were retained. The mean speed data obtained was limited to 200 samples per condition to plot.

### Measurements of sfGFP-TUA1 signal intensity

To quantify sfGFP-TUA1 signal intensity, cells were imaged as described above, and the acquired images were processed using ImageJ (National Institutes of Health). For analysis, a 50-µm-wide line was drawn along the region spanning from the apical to posterior ends of furrow-associated microtubules. The plot profile function in ImageJ was used to extract fluorescence intensity values along the selected regions, enabling quantitative comparison of TUA1-GFP signal distribution.

## Acknowledgments

We thank Dr. Ritsu Kamiya (Chuo University) for critically reading the manuscript and providing insightful comments. This work was supported by Takeda Science Foundation (to T.K.), Institute for Fermentation, Osaka (to T.K.), The Kato Memorial Bioscience Foundation (to T.K.), Research Grant from Human Frontier Science Program [RGP006/2023 (to T.O.) (https://doi.org/10.52044/HFSP.RGP0062023.pc.gr.168592)], Japan Society for the Promotion of Science [23K05829 (to T.K.), 21H02654 (to T.O.) and 24H02285 (to T.O.)], and the National Science Foundation MCB CAREER Award (2337141 to M.O.).

---

**Supplemental Figure 1.**
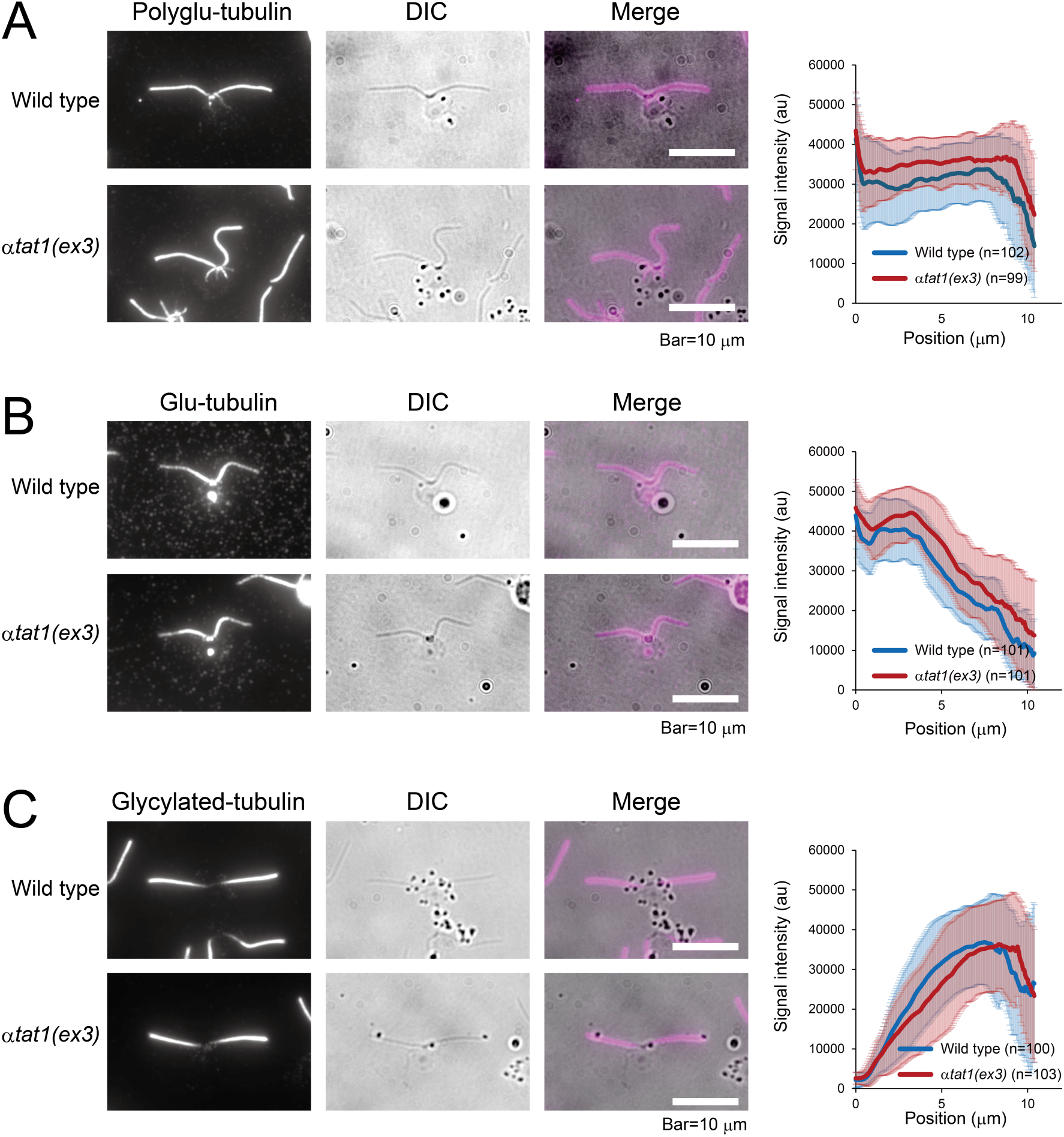
The levels of tubulin modifications in the *αtat1(ex3)* axoneme. Indirect Immunofluorescence microscopy of the NFAps isolated from wild type (cc124) and *αtat1(ex3)* using (A) anti-polyglutamylated tubulin antibody (polyE#2), (B) anti-glutamylated tubulin antibody (GT335), and (C) anti-glycylated tubulin antibody (Gly-pep1). Average signal intensities of the respective antibodies, measured along the length of the axoneme, are shown (right panels).

**Supplemental Table 1.**
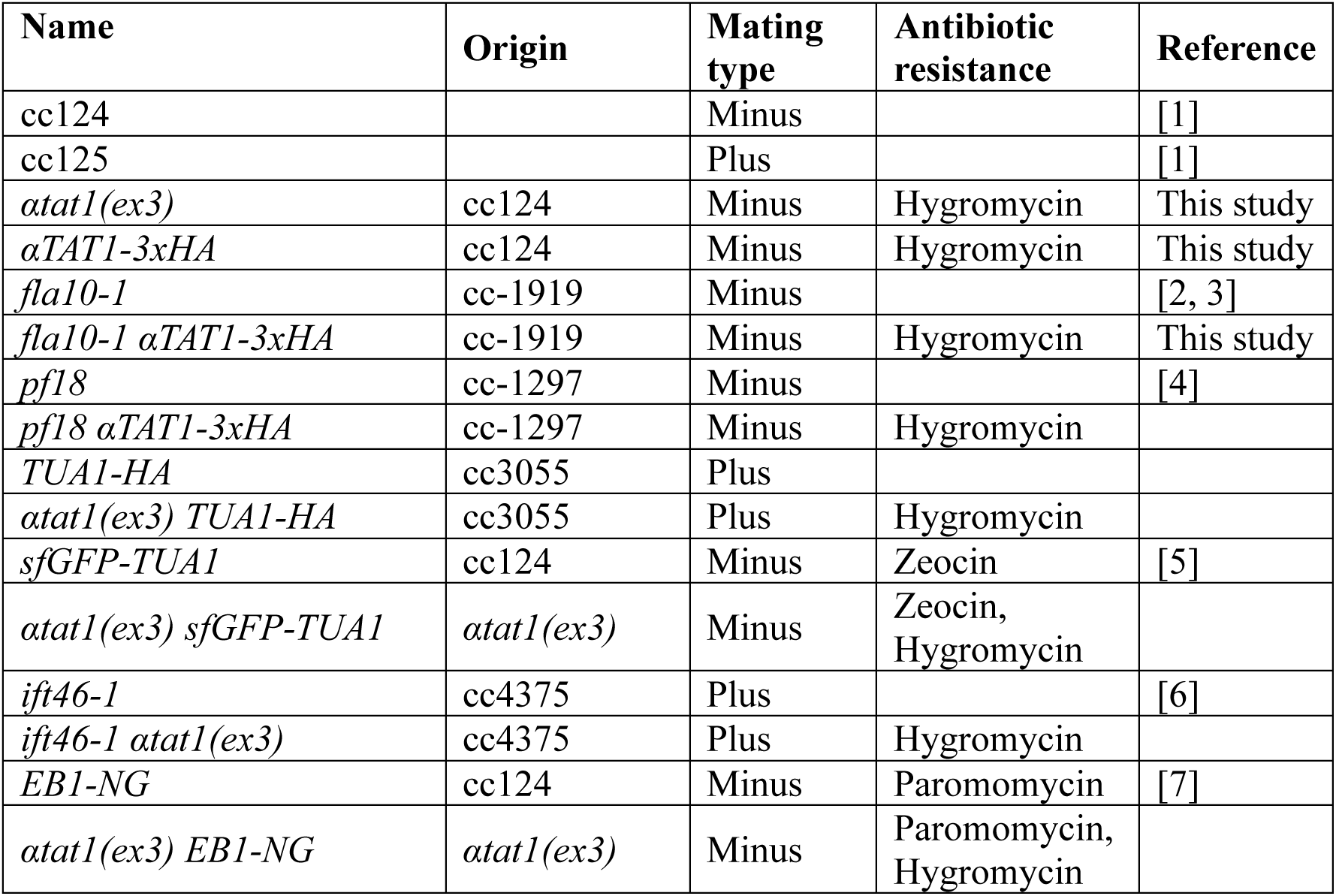
List of strains used in this study. *Generated mutants will be deposited to the *Chlamydomonas* Resource Center (https://www.chlamycollection.org/) after publication.

**Supplemental Table 2.**
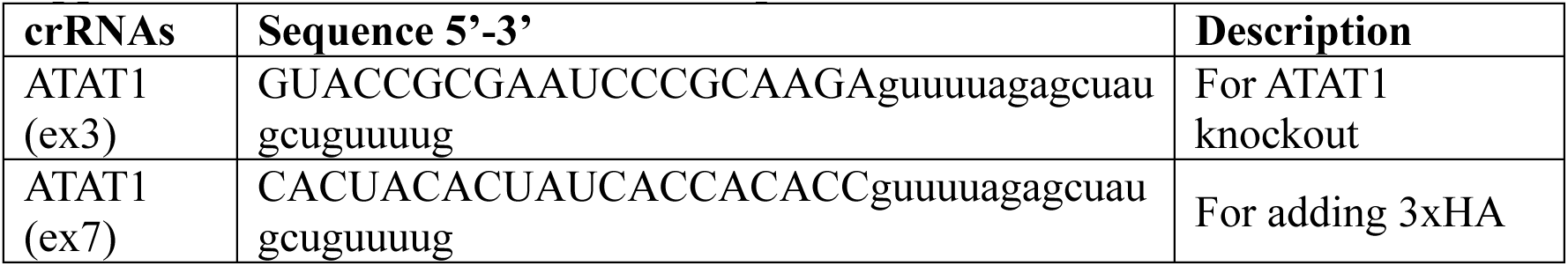
List of crRNA sequences.

**Supplemental Table 3.** Antibody used in this study.

| Antibody (clone) | Host | Dilution | Reference / Origin |
| --- | --- | --- | --- |
| Anti-acetylated tubulin (6-11B-1) | Mouse | 1:5000 (WB)<br>1:200 (IFM) | Sigma-Aldrich |
| Anti-tyrosinated tubulin (TUB-1A2) | Mouse | 1:3000 (WB)<br>1:200 (IFM) | Sigma-Aldrich |
| Anti-detyrosinated tubulin (AB3201) | Rabbit | 1:2000 (WB)<br>1:200 (IFM) | Merck |
| Anti-polyglutamylated tubulin (polyE#2) | Rabbit | 1:2000 (WB),<br>1:200 (IFM) | Kubo et al. (2017) |
| Anti-glutamylated tubulin (GT335) | Mouse | 1:2000 (WB),<br>1:200 (IFM) | Adipogen Life Sciences |
| Anti-glycylated tubulin (Gly-pep1) | Rabbit | 1:2000 (WB),<br>1:200 (IFM) | Funakoshi |
| Anti-α-tubulin (B512) | Mouse | 1:5000 (WB)<br>1:500 (IFM) | Sigma-Aldrich |
| Anti-HA (4B2) | Mouse | 1:2000 (WB) | FUJIFILM Wako Pure Chemical Corporation |
| Anti-HA (F10) | Rat | 1:200 (IFM) | Sigma-Aldrich |
| Anti-ATAT1N#1 | Rabbit | 1:2000 (WB) | This study |
| Anti-ATAT1N#2 | Rabbit | 1:2000 (WB) | This study |
| Anti-ATAT1C#1 | Rabbit | 1:2000 (WB) | This study |
| Anti-ATAT1C#2 | Rabbit | 1:2000 (WB) | This study |
| Anti-IFT172 | Mouse | 1:20 (WB) | DSHB |
| Anti-IFT139 | Mouse | 1:100 (WB) | DSHB |
| Anti-IFT81 | Mouse | 1:100 (WB) | DSHB |
| Anti-IFT57 | Mouse | 1:20 (WB) | DSHB |
| Anti-RPL4 | Rabbit | 1:2000 (WB) | This study |

**Supplemental Table 4.**
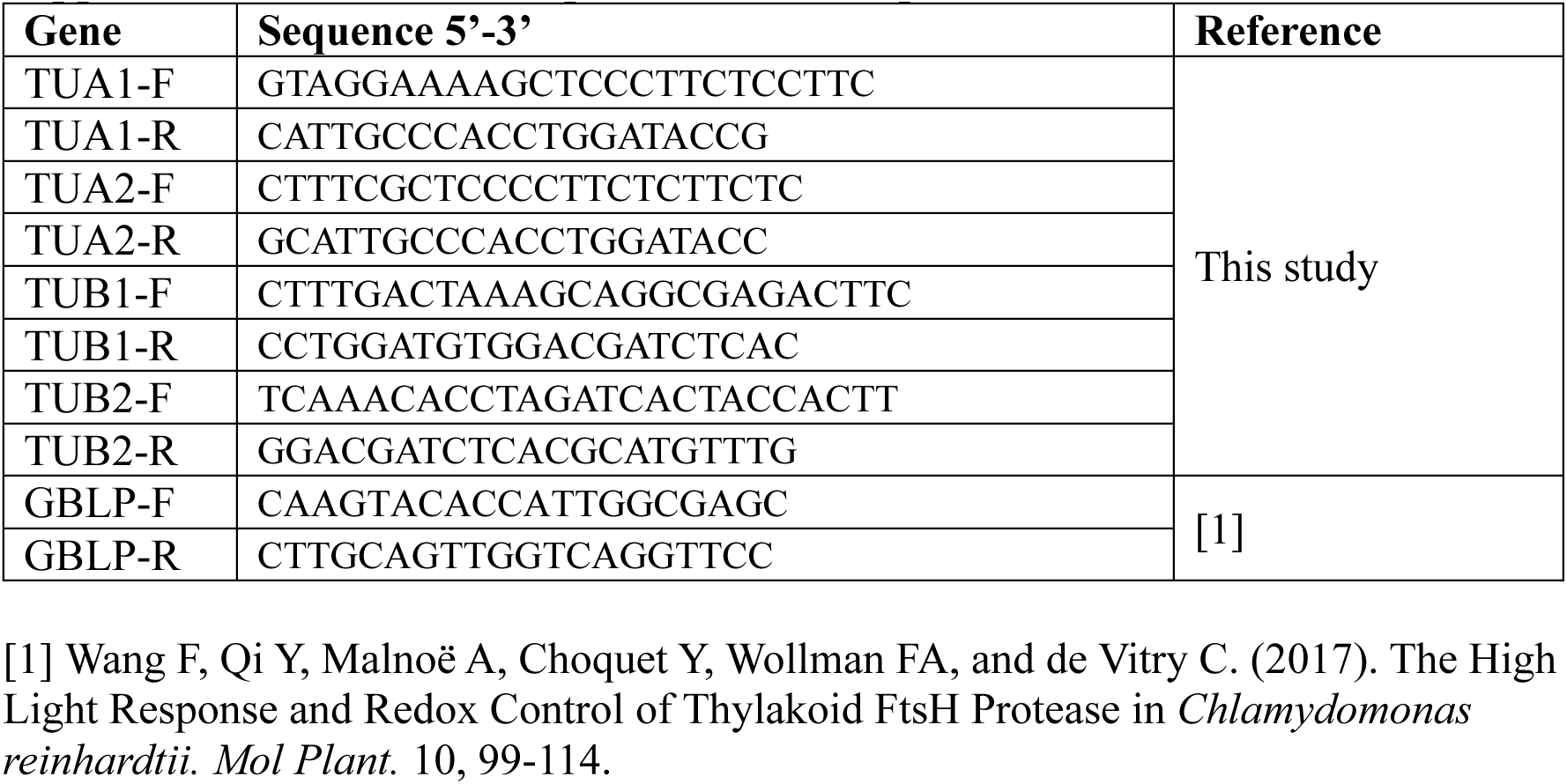
List of primers used for qPCR.

